# Intraventricular CXCL12 is neuroprotective and increases neurogenesis in a murine model of stroke

**DOI:** 10.1101/584565

**Authors:** Laura N. Zamproni, Marimelia A. Porcionatto

## Abstract

Stroke is the leading cause of physical disability and the second leading cause of death in adults. Chemokines that regulate ischemic microenvironment can influence neurorestorative therapies in stroke patients. CXCL12 was shown to be neuroprotective, but there are some contradictory results in the literature. The objective of this work is to verify whether CXCL12 delivery to the brain could be beneficial or harmful. We decided to evaluate the intraventricular delivery to facilitate its application in clinical practice. Intraventricular CXCL12 was able to produce increased mRNA expression of the receptors CXCR4 and CXCR7 in the cerebral cortex. After the stroke, intraventricular CXCL12 decreased mice ischemic area and improved behavioral response. We found CXCL12 was neuroprotective, increase reactive astrogliosis and improve neurogenesis in the perilesional area.

## Introduction

Stroke is the leading cause of physical disability and the second leading cause of death in adults (Machado et al. 2013). Although thrombolytic therapy and neuroendovascular intervention represents an advance for the treatment of acute stroke a large proportion of survivors still struggle with severe disabilities due to the limited time window for these therapies and frequent unsuccessful recanalization (Nakagomi et al. 2015). There is no definitive therapy that can restore lost brain function (Bang 2009). Numerous approaches have been developed to protect the brain from ischemic damage but have limited success in clinical practice (Bang 2009).

The microenvironment in the brain infarct area has been shown to influence the effects of neurorestorative therapies in stroke patients (Bang 2017). Regulating this microenvironment, by controlling chemokine and trophic factors levels, can increase the degree of recovery after stroke (Wang et al. 2007). Among the milieu of chemicals expressed in the injured tissue, C-X-C chemokine ligand 12 (CXCL12), also known as stromal-derived factor-1 (SDF-1) and its receptor CXCR4, are potential candidates to modulate the microenvironment in the infarcted brain (Shyu et al. 2008; Wang et al. 2012).

CXCL12 is a chemokine known to regulate migration, proliferation, and differentiation of neural stem cells (NSCs) within the developing central nervous system (CNS) (Li et al. 2015). Previous treatment of stroke with CXCL12 was shown to upregulates the CXCL12/CXCR4 axis and enhances neurogenesis, angiogenesis, and neurological outcome (Shyu et al. 2008; Yoo et al. 2012; Kim et al. 2015).

However, there is some contradictory results in the literature. CXCL12 was shown to function mainly as an inflammatory initiator during the acute phase of ischemia (Huang et al. 2013; Ruscher et al. 2013). Pharmacological inhibition of CXCL12 binding to its receptors during the first week after stroke reduced inflammation and neurodegeneration and promoted long-term recovery, without affecting the infarct size (Wu et al. 2017). Also, higher circulating CXCL12 levels were shown to be linked to poor outcome in stroke patients (Pan et al. 2016; Cheng et al. 2017a).

The objective of this work is to verify whether CXCL12 delivered to the brain could be beneficial or harmful to stroke outcome. Since chemokines must be delivered locally due to possible dangerous systemic effect, which severely limits their therapeutic use and all previous work used intraparenchymal delivery, we decide to evaluate the intraventricular delivery, in order to facilitate its application in clinical practice.

## Material and Methods

### Intraventricular injection

Seven-week C57BL/6 female mice were submitted to surgery under anesthesia with intraperitoneal administration of acepromazine (1 mg/Kg, Vetnil, Louveira, São Paulo), xylazine (10 mg/kg, Ceva, São Paulo, Brazil) and ketamine chloridrate (100 mg/kg, Syntec, São Paulo, Brazil). Peroperative analgesia was provided by fentanyl (0.05 mg/Kg, Cristália, São Paulo, Brazil). All procedures were approved by the Committee of Ethics in the Use of Animals (CEUA/Unifesp 9745220217). Intraventricular injection was performed according to a previously described protocol (DeVos and Miller 2013). Briefly, animals were placed under a stereotaxic frame, skin and skull were open. A 5 µL Hamilton syringe was used for injection (stereotaxic coordinates from bregma: AP +0.3 mm; ML +1.0 mm; DV −3.0 mm). Animals received 5 µL of PBS or CXCL12 20 ng/µL. Brains were collected 1 or 3 days later. Mice were anesthetized and euthanized by cervical dislocation. Ipsilateral brain cortex was immediately separated and frozen. Ipsilateral brain cortex to injection was used for RNA extraction.

### RNA extraction and Quantitative PCR (qPCR)

Total RNA was isolated from brain cortex using the Trizol® reagent (Life Technologies, Thermo Fisher, Waltham, EUA), and RNA concentrations were determined using a NanoDrop ND-1000 instrument (Thermo Fisher). Reverse-transcriptase reactions were performed with the ImProm-II Reverse Transcription System (Promega, Madison, USA) using 2 μg total RNA. qPCR was performed using Brilliant® II SYBR® Green QPCR Master Mix (Applied Biosystems, Thermo Fisher) and the Mx3000P QPCR System; MxPro qPCR software was used for the analysis (Stratagene, San Diego, CA USA). Primers sequences are shown in Table 1.

**Table 1:**
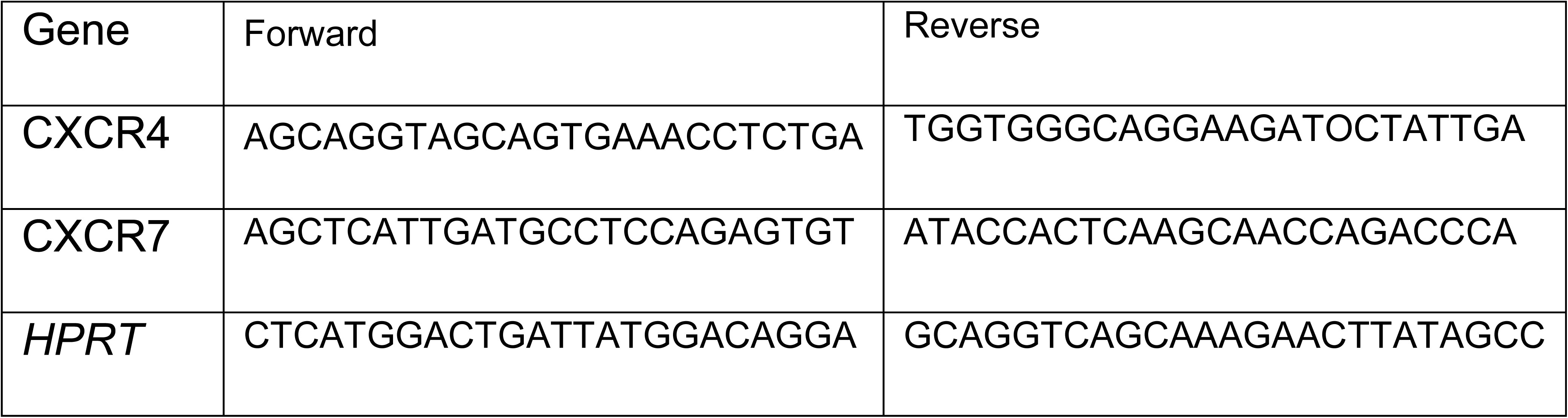
Primers sequences.

Values are expressed relative to those from the PBS group. For quantitation, target genes were normalized using a reference control gene (HPRT). The threshold cycle (Ct) for the target gene and Ct for the internal control were determined for each sample, and triplicate experiments were performed. The relative expression of mRNA was calculated using the 2^-ΔΔCt^ method (Livak and Schmittgen 2001).

### Stroke model

Seven-week C57BL/6 female mice were submitted to surgery under anesthesia with intraperitoneal administration of acepromazine (1 mg/Kg), xylazine (10 mg/kg) and ketamine chloridrate (100 mg/kg). Peroperative analgesia was provided by fentanyl (0.05 mg/ Kg). All procedures were approved by CEUA/Unifesp (9745220217).

Permanent distal middle cerebral artery occlusion was performed as previously described (Llovera et al. 2014). A 1 cm skin incision between the ear and eye was made and the temporal muscle was separated. The middle cerebral artery (MCA) was identified at the transparent skull, in the rostral part of the temporal area, dorsal to the retro-orbital sinus. The bone was thinned out with a drill right above the MCA branch and carefully withdraw.

MCA was coagulated with monopolar electrosurgery (MedCir, São Paulo, Brazil). The temporal muscle was relocated to its position, covering the burr hole and skin sutured. Immediately after, animals were placed under a stereotaxic frame, skin was open and skull was open. A 5 µL Hamilton syringe was used for intraventricular injection (stereotaxic coordinates from bregma: AP +0.3 mm; ML +1.0 mm; DV −3.0 mm). Animals received 5 µL of PBS or CXCL12 20 ng/ul immediately after stroke. Sham animals were submitted to whole procedure except artery coagulation. Mice were weighted every day. Brains were collected 7 days later. Mice were anesthetized and euthanized by cervical dislocation. Next, the brains were removed, stained with 1.5% TTC (2,3,5-Triphenyl-tetrazolium, Alphatec, São José dos Pinhais, Brazil) at room temperature, for 20 min and immersed in 4% PFA for 24 h. To estimate the amount of cerebral tissue lost after stroke, the brains were sectioned into 2 mm coronal slices using a manual sectioning block, and the lost tissue area in each slice was determined by subtracting the area of the injured hemisphere from the area of the normal hemisphere using Image J software. The volume of brain tissue lost was determined by the sum of each slice area multiplied by the thickness (2 mm): lost area = Σ (area of contralateral side – area of ipsilateral side) x 2 (Llovera et al. 2014). For histological analyses, brains were sectioned into 30 μm coronal slices at −20 °C using a CM 1850 cryostat (Leica, Wetzlar, Germany).

### Behavioral assessment

The functional outcomes of the stroke model were evaluated using two different behavioral tests, conducted one day before stroke and 1, 4, and 7 days after injury.

Cylinder: mice were recorded for 4 min while exploring an open-top, clear plastic cylinder. Forelimb activity while rearing against the wall of the cylinder is recorded. Forelimb use is defined by the placement of the whole palm on the wall of the cylinder, which indicates its use for body support. Forelimb use is expressed as a ratio of right/left-sided.

Grid walking: to verify limb sensitivity, mice were recorded while walking over a grid for 4 min. The brain injury was in the left cortex, so as the mouse walked over the grid, its right paws falls, because it could not grasp the grid properly. The number of falls reflects the severity of the lesion (López-Valdés et al. 2014).

### Fluoro-Jade B

Brain sections were mounted with distilled water onto gelatin coated slides and dried on a slide warmer at 37 °C. The tissue was fully dried within 20 min at which time it was immersed in 100% ethyl alcohol for 3 min followed by a 1 min change in 70% alcohol and a 1 min change in distilled water. The slides were then transferred to a solution of 0.06% potassium permanganate for 15 min. The slides were rinsed for 1 min in distilled water, 3 times, and were then transferred to the Fluoro-Jade (Sigma, St. Louis, EUA) staining solution for 30 min. A 0.01% stock solution of the dye was prepared by dissolving 10 mg Fluoro-Jade in 100 mL of distilled water. The 0.0001% working solution of Fluoro-Jade was prepared by adding 1 mL of the stock Fluoro-Jade solution to 99 mL of 0.1% acetic acid in distilled water. After staining, the sections were rinsed 1 min, 3 times, in distilled water. Excess water was drained off and the slides were immersed in xylene and then coversliped with DPX (Sigma) mounting media. Sections were examined with a scanning confocal inverted microscope (TCS, SP8 Confocal Microscope, Leica).

### Immunofluorescence

For immunofluorescence staining, cortical sections were incubated overnight at 4 °C with anti-glial fibrillary acidic protein (GFAP, 1:1000, chicken IgG, Merck Millipore, Burlington, EUA), anti-Iba1 (1:200, goat IgG, Abcam, Cambridge, United Kingdom), anti-doublecortin (DCX, 1:500, guinea-pig IgG, Merck Millipore) and anti-CD31 (1:200, BD, Franklin Lakes, USA). After washing with PBS, the sections were incubated at room temperature with the appropriate secondary antibodies conjugated to Alexa Fluor® 488 or 594 (1:500, Invitrogen, Carlsbad, USA). Nuclei were stained with DAPI (1:500, Molecular Probes, Eugene, USA). Glass slides were mounted using Fluoromount G (Sigma). The fluorescently labeled tissue slices were analyzed using a scanning confocal inverted microscope (TCS, SP8 Confocal Microscope, Leica), and image overlays were generated using ImageJ software. Stained cells were quantified by measurement of the total corrected fluorescence (Total corrected fluorescence = Integrated Density – (Area of selection X Mean fluorescence of background readings) (McCloy et al. 2014) or by manual cell counting per mm^2^. Data from 3 animals in each group, 3 sections per animal were analyzed.

### Statistical analysis

Statistically significant differences were evaluated by Student’s T-test or one-way ANOVA followed by Tukey post-test using the GraphPad Prism software version 5.01 (GraphPad Software, USA). Results are expressed as mean ± SEM and were considered significant if p<0.05.

## Results

### Intraventricular administration of CXCL12 increased CXCR4 and CXCR7 expression in the cerebral cortex

We first evaluated whether CXCL12 intraventricular injection could promote a response in the brain cortex (Figure 1). We found that animals who received CXCL12 presented an increase in CXCR4 expression after 24 h (Figure 1A) and significant increase in CXCR4 expression after 72 h (Figure 1B). CXCR7 expression significantly increased in the first 24 h (Figure 1C) and was similar to PBS after 72 h (Figure 1B).

**Figure 1:**
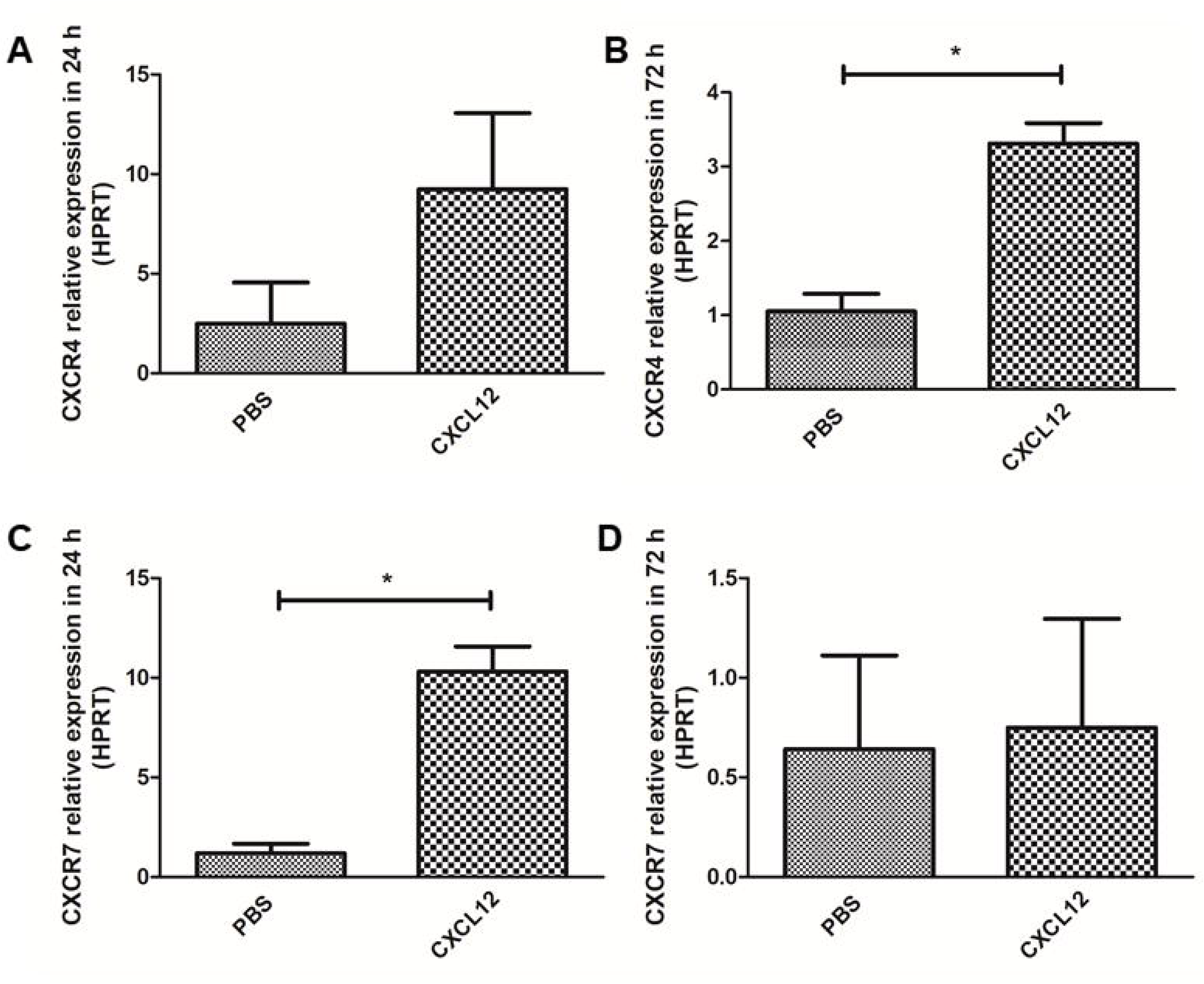
CXCR4 and CXCR7 expression in brain cortex after intraventricular CXCL12 administration. (A and B) CXCR4 relative expression in the cortex in 24 h and 72 h (*= p < 0,05, Student’s T-test, n=3). (C and D) CXCR7 relative expression in the cortex in 24 h and 72 h (*= p < 0,05, Student’s T-test, n=3).

### Intraventricular CXCL12 reduced infarction volume

Once we confirmed intraventricular delivery produced a cortical response regarding expression of CXCL12 receptors CXCR4 and 7, we evaluated its effect on stroke. We first found that mice receiving CXCL12 have a lower ischemic area confirmed by TTC staining (Figure 2A and B). Also, animals who received CXCL12 behave better, losing less weight after stroke, in a similar way with Sham group (Figure 2C). Although the cylinder test revealed no differences between the groups (Figure 2D), in the 7^th^ day post ischemia, animals who received CXCL12 recovery better, presenting fewer falls when walking in the grid (Figure 2E).

**Figure 2:**
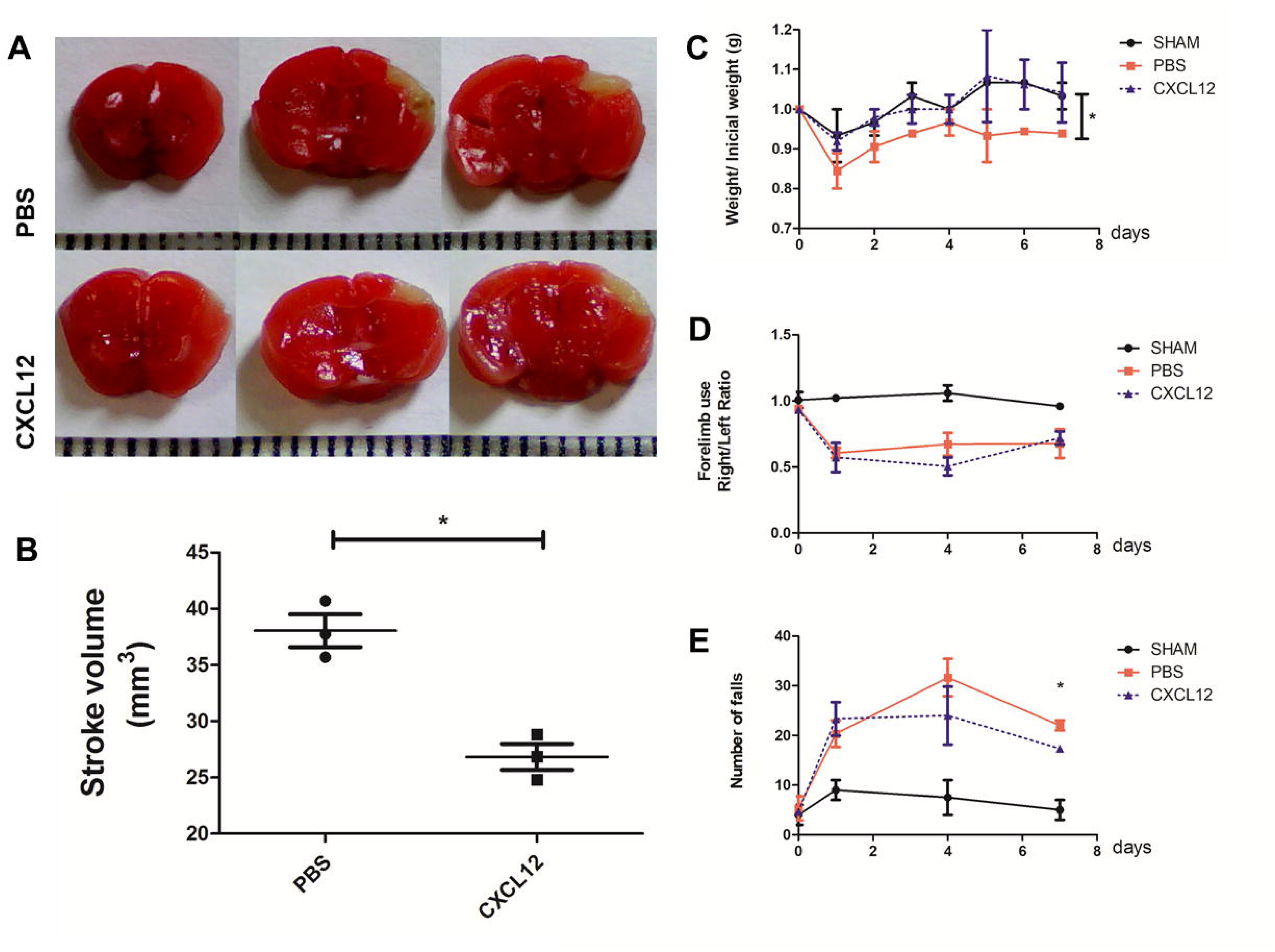
Impact of intraventricular CXCL12 on ischemic brain cortex. (A) TTC staining of brain slices. Unstained area corresponds to death tissue. (B) Quantification of stroke area by TTC method (*= p < 0,05, Student’s T-test, n=3). (C) Weight loss/gain curve (*= p < 0,05 for SHAM x PBS, CXCL12 x PBS, one-way Anova plus Tukey posttest n=3). (D) Right/left ratio for forelimb use in the cylinder test (n=3). (E) Number of paws falls in the grid walking test (*= p < 0,05 for SHAM x PBS, SHAM x CXCL 12, one-way Anova plus Tukey posttest n=3). TTC: triphenyltetrazolium chloride

### Intraventricular CXCL12 decreased neuronal degeneration and increased reactive astrocytes in brain ischemia

Our next step was to evaluate the molecular mechanisms involved in stroke response to CXCL12 administration.

We first evaluated neuroprotection by analyzing the presence of degenerating neurons, stained by Fluorojade B, in the ischemic borders (Figure 3A). Mice who received CXCL12 presented significantly less neuronal degeneration (Figure 3B).

**Figure 3:**
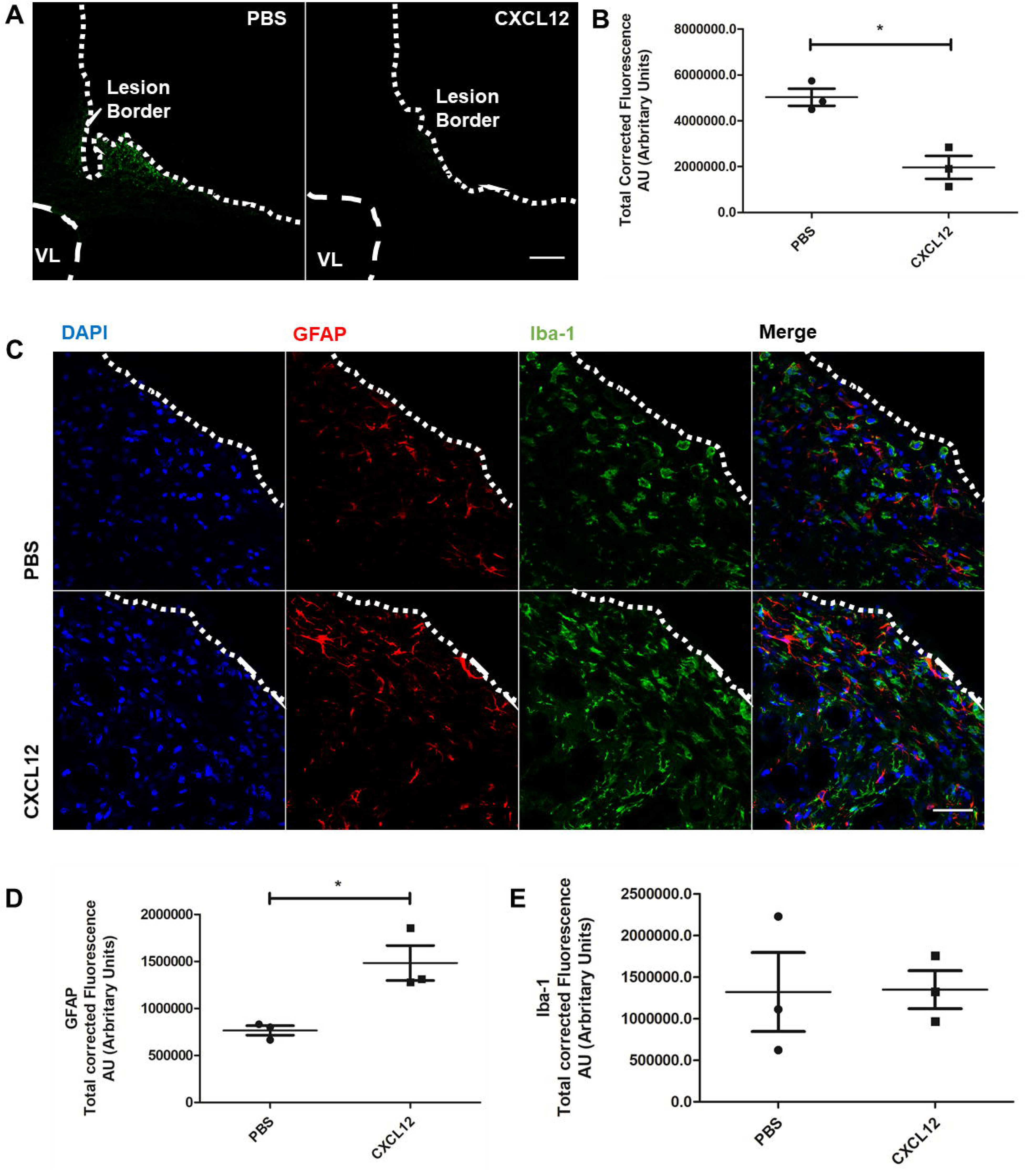
CXCL12 promotes neuroprotection and increases astroglial response in the ischemic brain cortex. (A) Fluoro-Jade B staining for neuronal degeneration (Scale bar =200 µm). (B) Total corrected fluorescence for Fluoro-Jade B staining (*= p < 0,05, Student’s t-test, n=3). (C) Microglial and astrocyte lesion infiltrate in 7 days. Immunohistochemistry to IBA-1 (microglia) ang GFAP (astrocytes) (scale bar = 50 µm). (D) Total corrected fluorescence for GFAP staining (*= p < 0,05, Student’s t-test, n=3). (E) Total corrected fluorescence for IBA-1 staining (n=3). LV: lateral ventricle; AU: arbitrary units. DAPI: 4′,6-diamidino-2-phenylindole; IBA-1: Ionized calcium binding adaptor molecule 1; GFAP: *Glial fibrillary acidic protein.*

Surprisingly, along with less neuronal degeneration we observed an increase of the GFAP infiltrate around the ischemic area in 7 days (Figures 3C and D). The microglial infiltrate, however, did not change between PBS and CXCL 12 groups (Figures 3C and E).

### Intraventricular CXCL12 increased neurogenesis but not angiogenesis

Finally, we evaluated neurogenesis and angiogenesis at the lesion site. We found twice as many neuroblasts in CXCL12 treated animals (Figures 4A and B), but we did not observe an increase in the number of endothelial cells (Figures 4C and D).

**Figure 4:**
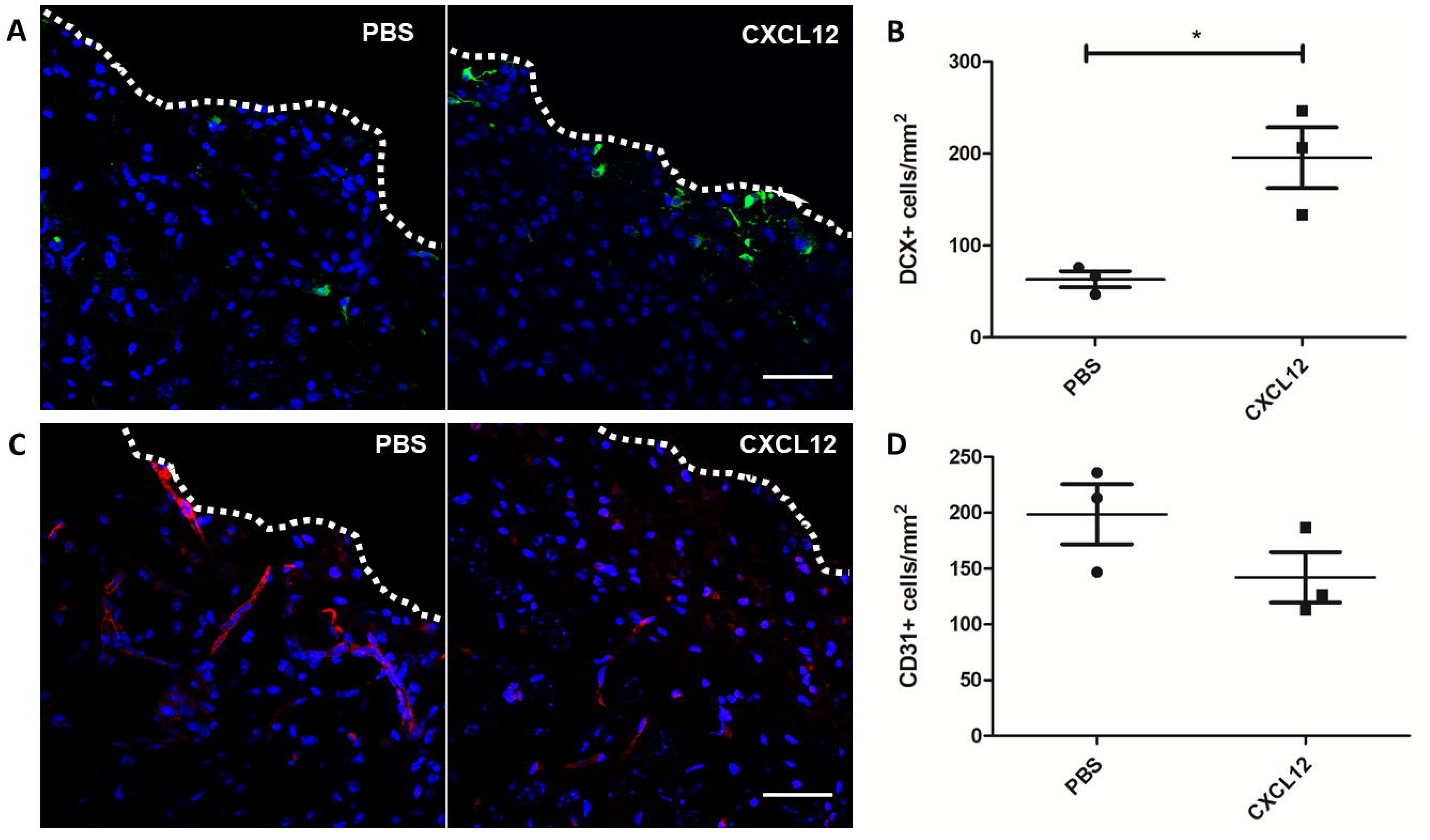
CXCL12 increases neurogenesis in ischemic brain cortex. (A) Immunohistochemistry to DCX (neuroblasts) (scale bar = 50 µm). (B) Number of DCX stained cells per mm^2^ (*= p < 0,05, Student’s t-test, n=3). (C) Immunohistochemistry to CD31 (endothelial cells) (scale bar = 50 µm). (D) Number of CD31 stained cells per mm^2^ (n=3). DCX: doublecortin, CD: cluster of differentiation.

## Discussion

We chose an intraventricular injection to deliver CXCL12 because stroke is usually an extensive lesion and we wanted CXCL12 to reach all the cortex. If we performed an intraparenchymal injection, CXCL12 would concentrate in a single region and probably not diffuse to the entire lesion. We could overcome this by performing multiple intraparenchymal injections, but then, it would be very difficult to translate the results into clinical practice, due the morbidity induced by the procedure.

Our data support that CXCL12 delivered by intraventricular via was able to increase expression of CXCL12 receptors in brain cortex and exert the neuroprotective effects observed previously in brain ischemia (Shyu et al. 2008; Yoo et al. 2012; Kim et al. 2015). CXCR4 is found at the membrane of a variety of cells including neurons, astrocytes, microglia, bone marrow-derived cells, and NSCs (Wang et al. 2012). CXCL12 and CXCR4 are constitutively expressed in the brain but are up-regulated in the ischemic penumbra regions following ischemic stroke (Cheng et al. 2017b). CXCL12/CXCR4 is thought to play important roles in multiple processes after ischemic stroke, which include inflammatory response, angiogenesis, and the recruitment of bone marrow-stem cells and NSCs to injury (Kokovay et al. 2010; Yellowley 2013). Despite this evidence, there is plenty of data in the literature showing that AMD3100, a CXCR4 inhibitor, can improve outcome after stroke, mainly by attenuating inflammatory response (Huang et al. 2013; Ruscher et al. 2013; Wu et al. 2017). However, AMD3100 can also bind CXCR7 and function as an allosteric agonist (Kalatskaya et al. 2009).

CXCR7 affinity for CXCL12 is almost 10 times higher than for CXCR4 and is more distributed in neurons (Cheng et al. 2017b). The pathways behind their role are not completely understood. CXCR7 was thought to function as a scavenger receptor that can regulate the extracellular availability of CXCL12, mediating its internalization and subsequent lysosomal degradation (Boldajipour et al. 2008; Sánchez-Alcañiz et al. 2011). Clarification of the role of CXCR7 will be required in the future (Cheng et al. 2017b). However, lack of a commercial CXCR7 antagonist difficult this process.

We found that intraventricular CXCL12 decreased degenerating neurons along ischemic area. In fact, in brain development, CXCL12 was find to control neuronal surviving and inhibiting apoptosis (Khan et al. 2008). Another interesting finding was the increased number of reactive astrocytes in the perilesional area. Astrocytes are commonly thought to be involved in negative responses through glial scar formation and inhibition of axonal regeneration (Silver et al. 2014). But new data describe that astrocytes appear to possess a high potential for regeneration and neuroprotection following stroke (Sofroniew and Vinters 2010; Becerra-Calixto and Cardona-Gómez 2017; Teh et al. 2017). Astrocytes are involved in a large number of key processes essentials to keep integrity of the nervous system, such as regulating the vascular tone, removing excess glutamate in synaptic cleft and preventing excitotoxicity, releasing neurotrophic factors and antioxidants and promoting synaptogenesis (López-Valdés et al. 2014). Many studies have suggested that after cerebral ischemia the astrocytes are able to de-differentiate into NSCs and provide a new source of neurons at the lesion site (Magnusson and Frisén 2016). To our knowledge, the increase in reactive astrocytes in response to CXCL12 have not been reported.

Our most disappointing data was the lack of increased angiogenesis in response to CXCL12 in contrast to what was previously described in the literature. One possible explanation for this is that most studies evaluated angiogenesis after a longer time period, usually 2 to 5 weeks, whereas we used a 7-day end-point (Shyu et al. 2008; Li et al. 2018).

## Conclusions

Intraventricular administration of CXCL12 was able to produce increased mRNA expression of the receptors CXCR4 and CXCR7 in the cerebral cortex. In stroke, intraventricular administration of CXCL12 decreased mice ischemic area and improved behavioral response. We found CXCL12 was neuroprotective, increased reactive astrogliosis and improved neurogenesis in stroke area.

## Acknowledgments

We kindly thank FAPESP (MP: 2009/05700-5, 2012/00652-5) and CNPq (LNZ: 380304/2017-1, MP: 404646/2012- 3, 465656/2014-5) for financial support.

